# Learning may help pollinators find their host plants in polluted landscapes

**DOI:** 10.1101/649251

**Authors:** Brynn Cook, Alexander Haverkamp, Bill S. Hansson, T’ai Roulston, Manuel Lerdau, Markus Knaden

## Abstract

Pollination strongly contributes to food production, and often relies on pollinating insects. However, atmospheric pollution may interfere with pollination by disrupting floral plumes that pollinators use to navigate to flowers.

In this study, we examine the impacts of pollution-induced elevated ozone levels on the composition of a floral blend of *Nicotiana alata* and examine the response of innate and trained *Manduca sexta* to the ozone-altered blend.

Ozone exposure altered the floral blend of *N. alata*, and disrupted the innate attraction of naïve *M. sexta* to the altered blend. However, associative learning can offset this disruption in attraction. Moths that were enticed with visual cues to an artificial flower emitting an ozonated blend learned to associate this blend with a nectar reward after just one rewarded experience. More importantly, moths that were rewarded while experiencing the unozonated floral blend of their host subsequently found the ozonated floral blend of the same host attractive, most likely due to experience-based reinforcement of ozone-insensitive cues in the blend.

The attraction of moths to both unaltered and ozonated plumes is critical for tolerating polluted landscapes. At the host plant, where moths feed, floral emissions are relatively pure. As floral odors travel away from the host, however, they become degraded by pollution. Therefore, targeting the flower requires recognizing both conditions of the odor. The ability to generalize between the pure and ozone-altered scents may enable pollinators like *M. sexta* to maintain communication with their flowers and reduce the impact anthropogenic oxidants may have on plant-pollinator systems.

## 1 INTRODUCTION

Pollination is integral to maintaining diverse and healthy ecosystems (Kevan 1999), and it strongly contributes to global food production (Klein et al 2007). The coevolutionary relationship between plants and their pollinators is maintained when plants emit signals that pollinating insects can detect and recognize as belonging to a host plant. These signals include visual cues, such as brightly colored flowers, and olfactory cues – i.e. floral scents (Kunze and Gumbert 2001). Because the visual acuity of insect pollinators is limited to a resolution of centimeters to a few meters for most flowers (Kapustjansky et al. 2010, but see Ohashi and Yahara (2001) for visual detection of flower patches over longer distances), smell is likely to be an important sensory modality guiding pollinators to flowers over long distances. Floral scents are comprised of bouquets of volatile organic compounds (VOCs) that are emitted from a flower and travel downwind, forming a “scent pathway” that can lead pollinators through the landscape to their host plant.

Animal pollination depends on plants producing signals that are maintained in the landscape and that pollinators can recognize, yet the plume composition of a given plant species may vary both spatially and temporally. Alterations in floral scent over evolutionary time scales, such as changes in scent due to modifcations in genes coding for specific VOCs, have been shaped by the coevolutionary partnership between plants and pollinators (Dudareva and Pickersky 2000). However, variation in plume production can also occur outside of the prescriptions of plant-pollinator coevolution, including variations at sub-evolutionary timescales, and a pollinator will encounter variable plumes over its lifetime, even if it forages on just one plant species. In unpolluted environments, floral plumes can vary due to differences both in emissions and in changes in the plume that occur post-emission as it moves through the atmosphere. A plant’s emission of floral plumes can vary over time, due to both diel cycles (e.g. Theis et al. 2007; Raguso et al 2003; Muhlemann 2014) and seasonal cycles as plants progress through different phenological states (e.g. Theis et al 2007; Desurmont et al 2015). Furthermore, production of plumes may vary across space: a population of one species in the landscape may have a different genetic expression for floral scent than another patch of the same species (Knudsen et al. 2006, Haverkamp et al. 2018), and environmental gradients such as soil moisture and nutrient load can also influence plume production (Majestic et al. 2009). Post-emission plume transformation also contributes to the variability of floral scents across a landscape: changes to wind speed, temperature, and turbulence all affect the concentration of a plume that the insect experiences and the probability of and frequency at which a pollinator encounters a plume (Murlis et al 1992; Finelli et al 2000). Likewise, a pollinator’s location in the landscape and its distance from the emitting host plant will dictate the frequency with which it encounters a floral scent (Visser et al 1986).

Given the variability in floral scents within the spatial and temporal foraging breadth of a single insect, pollinators must exhibit strategies to cope with plume variation. Pollinators could simply manifest a broad innate attraction to many floral compounds (e.g., Bisch-Knaden et al 2018), such that a relatively stable subset of compounds provides a reliable cue in the midst of variation. If a broad innate recognition of cues is not sufficient to maintain attraction to variable floral scents, pollinators can use another coping strategy: learning. Pollinators can modulate their preference for flowers based on their experience, so that an insect with no innate recognition for a specific floral compound or blend of compounds can associate that olfactory signal with a nectar reward while feeding at the flower (Wright and Schiestl 2009). Learning in this way leads to an increased repertoire of olfactory cues; hawkmoths readily forage on an innately less-preferred host plant after olfactory-association, but they maintain their innate recognition for preferred host flowers (Riffell et al 2013). Learning may also assist pollinators in recognizing floral blends that have been modified as they move downwind of the plant; after learning an odor, honeybees can later recognize that same odor at a lower concentration (Bhagavan and Smith 1997; Pelz et al 1997). Another type of learning even enables insects to recognize compounds they have no first-hand associative experience with; pollinators may generalize their experience with one compound to another compound, if both compounds have a similar chemical structure (Daly et al 2001). By responding similarly to chemically related compounds, pollinators can follow plumes that differ at production due to plant genetics or physiological conditions, or that differ because of modest chemical shifts in the plume as it travels away from the plant. In these ways both plasticity in behavior and innate recognition of compounds may work together to enable pollinators to cope with variable olfactory cues.

The conditions causing plume variation described above have existed over evolutionary time scales, and thus the composition and concentration of floral plumes produced by plants has been subjected to selection pressures within the overall coevolutionary relationship between plant species and their primary pollinators (e.g., Dudareva and Pichersky 2000). In this manner, the ability of pollinators to find rewarding host plants despite variation in chemical signals has been subject to selection pressure. Today, however, many pollinators face landscapes with steeply increased floral plume variation as a result of anthropogenic interferences (Jurgens and Bischoff 2017).Following the industrial revolution, there has been a dramatic increase in the tropospheric load of atmospheric pollutants, including the oxidant species nitrate radical and ozone, and potentially hydroxyl radical (Naik et al 2013; Spivakosky et al 2000; Hauglustaine and Brasseur 2001). All three tropospheric oxidants are strongly reactive and can break apart components of floral blends. All three of these highly reactive oxidants can react with the carbon-carbon double bonds that are commonly found in floral volatiles (Atkinson and Arey 2003; Baker et al 2002). Because floral volatiles have different structures and thus react differently from each other with a given oxidant, oxidant pollution leads to both decreases in some key compounds and changes in the relative concentrations of individual compounds in a floral blend (Lusebrink et al 2015; McFrederick et al 2008; Farre-Armengol et al 2016) – both of which may be important cues to foraging insects (Bruce et al 2005). Moreover, when oxidants react with a VOC, they induce a series of reactions that can lead to the production of secondary compounds. Secondary compounds may themselves be long-lived VOCs, many of which are compounds that share little similarity to the parent VOCs, e.g. formaldehyde, acetone, and carbon monoxide. (McFrederick et al 2008; Lee et al 2006). While previous work finds that a pollinator can cope with ‘noise’ in a floral blend resulting from addition of non-target biological VOCs as well as anthropogenic VOCs (Riffell et al 2014), it is unclear if pollinators can maintain attraction to plumes that lose compounds at different rates due to oxidation.

Tropospheric oxidants thus have the potential to alter floral scent trails and impede pollinators attempting to locate host plants. One oxidant, ozone has increased from approximately 10ppbv or less in preindustrial times (Hauglustaine and Brasseur 2001) to current averages in North America of 20-45ppbv (Vingarzan 2004), with spikes as high as 120ppbv during summertime ozone events (Fiore 2002; Vingarzan 2004). To continue using floral scents as cues in a world with elevated tropospheric ozone, pollinators must either hone in on non-reactive volatile compounds, or they must learn the succession of odors they encounter in the landscape, ranging from highly ozone-altered blends at distances far from the host plant, to blends that are unaltered at the flower. Current work has established that ozone-altered floral blends are less attractive than unaltered blends to a variety of insects including a bumblebee and two specialist herbivores (Farre-Armengol 2016; Fuentes et al 2013; Li et al 2016). Chemical modelling studies have predicted that ozone will react with floral blends across landscapes (McFrederick et al 2008 & 2009) and that as a result, insects will be less adept at locating their host plants in ozone-enriched environments (Fuentes et al 2016). While these works have demonstrated the potential for ozone to alter floral blends and impede insect foraging, neither empirical nor computational studies have considered the ability of insects to learn to identify and respond to ozone-altered blends. Can the flexibility in cue recognition or the learning abilities of pollinators, established to help these insects thrive in variable landscapes, assist them in recognizing floral cues even as air pollution alters the integrity of those blends? We test the ability of one nighttime pollinator, the hawkmoth *Manudca sexta*, to recognize and learn ozone-altered floral blends of one of its preferred host flowers, *Nicotiana alata*, using ozone as a substitute for the nighttime dominant nitrate radical; although the specific reaction chemistry of nitrate radical and ozone are distinct, both have the potential to alter floral blend composition via reaction with floral volatiles. Moreoever, while ozone is much more abundant in the troposphere than nitrate radical, it is also much less reactive, so that the lifetime of floral volatiles with ozone in the troposphere is typically longer than the lifetime of floral volatiles with nitrate radical, making ozone a conservative substitute for nitrate radical. After demonstrating that ozone substantially alters the odor profile of *N. alata* and renders it unattractive to naive moths, we develop an odor learning protocol for *M. sexta,* and consider two possible learning scenarios in which *M. sexta* could navigate to its host using odor cues, despite the alteration of the floral blend by ozone. First, we test whether *M. sexta* can can learn to use an ozonated blend as a cue that leads to a pure blend at a rewarding artificial flower. Second, we test whether experience at an unpolluted host broadens the suite of cues used in host recognition such that the floral blend remains attractive despite ozonation.

## 2 MATERIALS and METHODS

### 2.1 Study Organisms Rearing

*Manduca sexta* were raised in a light, humidity, and temperature controlled chamber (light:dark = 16:8, 70% relative humidity and 25° C during the light phase, and 60% relative humidity and 20 °C during the dark phase). Chamber day and night were inverted so that the moths experienced nighttime conditions during the day and were active during normal working hours of the researchers. *Nicotiana alata* flowers used for odor collection and for behavior tests in the wind tunnel were reared in a temperature and light-controlled chamber with the same light and temperature specifications used for the *M. sexta* rearing chamber.

### 2.2 Effect of increased ozone levels on floral blend chemistry

We examined the effects of ozone on the blend composition of *N. alata* by comparing the floral blend mixed with scrubbed air to the blend mixed with air enriched with 120 ppbv ozone. Here, ozone serves as a conservative proxy for the nighttime dominant nitrate radical, as both ozone and nitrate can break apart carbon-carbon double bonds, but ozone is typically less reactive than nitrate radical. We placed two one-day old *N. alata* flowers into a small (200 ml) chamber. Air that had been run through a charcoal scrubber was blown at a rate of approximately 3 l/min into a chamber which contained two *N. alata* flowers. The floral headspace was then pulled from the chamber and split into two blends via two pumps, each pulling a little over 1 l/min. Both of the split floral headspaces were run through a flow meter and then into an airtight 2 l glass bottle at a rate of ~1 l/min. In one glass bottle, air (which had been scrubbed clean by being passed through a cylinder packed with glasswool and activated carbon pellets) flowed into a bottle containing half of the floral headspace at a rate of 0.5 l/min. In the other glass bottle, 0.5 l/min of ozone enriched air at a concentration just above 120 ppbv was mixed with the floral headspace: this concentration was set so that the ozone concentration in the mixed blend entering the wind tunnel was ~120 ppbv. Ozone used in these experiments was generated using a Thermo Scientific Ozone Generator (Model 165 Thermo Scientific in., Pittbsurg PA) and ozone concentrations were measured using an ozone analyzer (Model 202, 2B Technologies INc., Boulder CO). Both the blend mixed with scrubbed air and the blend mixed with ozone-enriched air (120 ppbv) were further homogenized while being pushed through an additional series of two 1 l bottles, before passing through a small box containing five 5 mm polydimethysiloxane (PDMS) tubes that captured the floral blends as they passed over them. This collection allowed direct comparison between the blend mixed with air and mixed with ozone-enriched air from the same flowers over the same mixing time.

Following the scent collection, PDMS tubes were analyzed individually using a thermo desorption unit (TDU, Gerstel, Germany) coupled to a temperature-programmable vaporizing unit (CIS 4, Gerstel, Germany), which was linked to an Agilent 7890A gas chromatograph (Agilent Technologies, CA) running in splitless mode and being connected to an Agilent 5975C mass spectrometer (electron impact mode, 70eV, ion source: 230◦C, quadrupole: 150°C,mass scan range: 33–350u) (Fig. 1A). We used a nonpolar column (HP-5 MS UI, 30m length, 0.25mm ID, 0.25µm film thickness, J and W Scientific, USA) under constant helium flow of 1.1 ml/min. The TDU temperature climbed from 30°C to 200°C at a rate of 100°C/min and held for 5min. Volatized compounds were trapped within the CIS 4 cooled injection system at −50°C and subsequently injected into the GC. The GC oven was programmed to hold 40°C for 3min, to increase the temperature at 5°C/min to 200°C, then to increase temperature at 20°C/min to 260°C, which was kept for 15min. Unprocessed data files were then exported and analyzed using the software package XCMS (Smith et al., 2006) implemented in R (R Core Team, 2014). Peak area values were log transformed to ensure normality and finally compared by a principal component analysis (Fig. 1B).

**Fig. 1.**
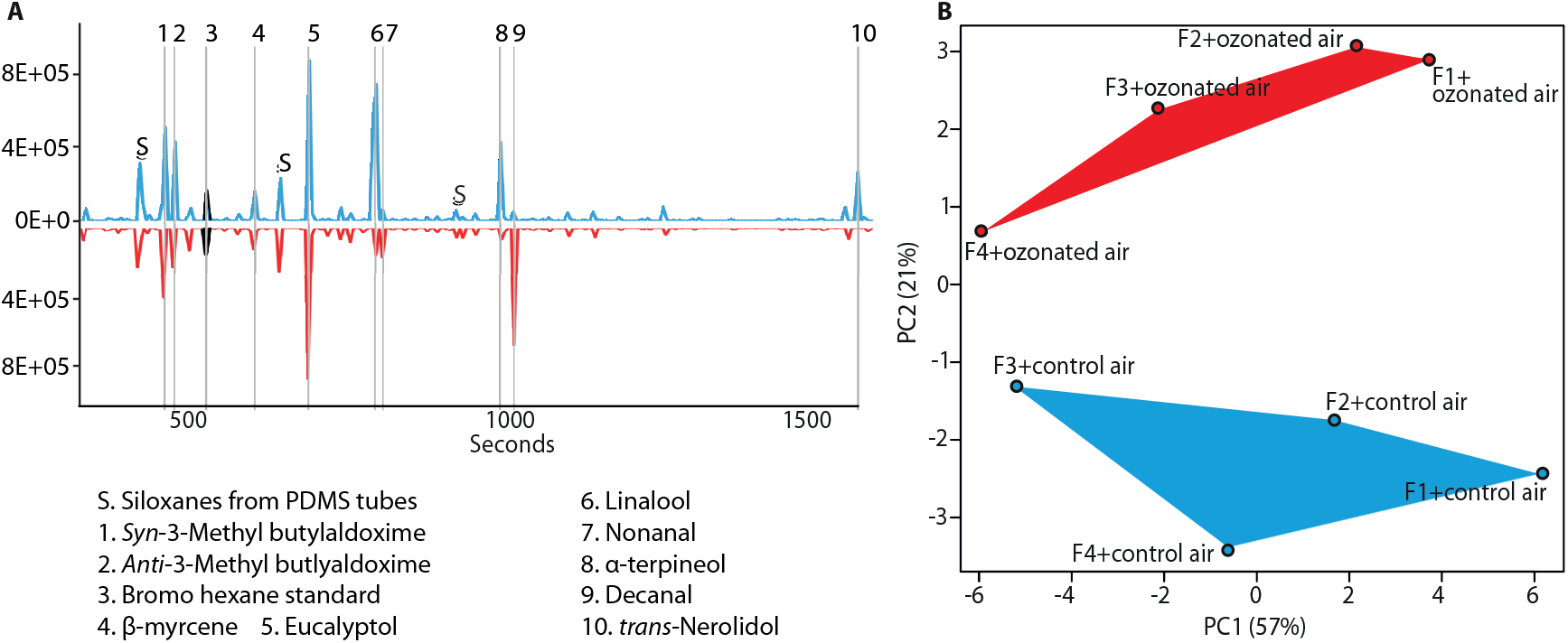
Floral blends of *N. alata* are altered by exposure to ozone. A. Example traces of original (blue) and ozonated (red) split headspaces of two *N. alata* flowers. Numbered peaks are identified by the NIST library (R-match > 90%). B. Principal component analysis of original and ozonated headspaces of four *N. alata* flowers (F1-F4).

### 2.3 Behavioral Assay Protocol

For all behavior assays, three-day old naïve *M. sexta* adults were collected from their climate chamber in individual mesh containers and given at least one hour to acclimate to the conditions in the wind tunnel in an adjacent room. The wind tunnel (airflow of 0.4 m s 1, 0.5 lux light, 25 °C and 70% humidity) itself was a plexiglas structure with dimensions L × H × W: 250 × 90 × 90 cm with dark circles randomly scattered on the tunnel floor that give visual feedback for flight stabilization (Fig. 2A). To start each behavior assay, a moth was placed inside the wind tunnel perched on the lid of its mesh container on a stand at the downwind side of the wind tunnel. Moths were mildly provoked to initiate flight by shaking the lid of the mesh container on which they were standing. Once flight was initiated, the moths were given four minutes to complete their behavior assay, during which time they were filmed by a series of cameras inside and outside of the wind tunnel. Scents were brought into the wind tunnel via Teflon tubing that was passed into upright metal cylinders. In some tests, paper artificial flowers were added atop these upright metal cylinders, and in others the visual cues were minimized and no artificial flowers were presented. The amount of time that the moths spent investigating a scent source with their extended proboscis was recorded manually.

**Fig 2.**
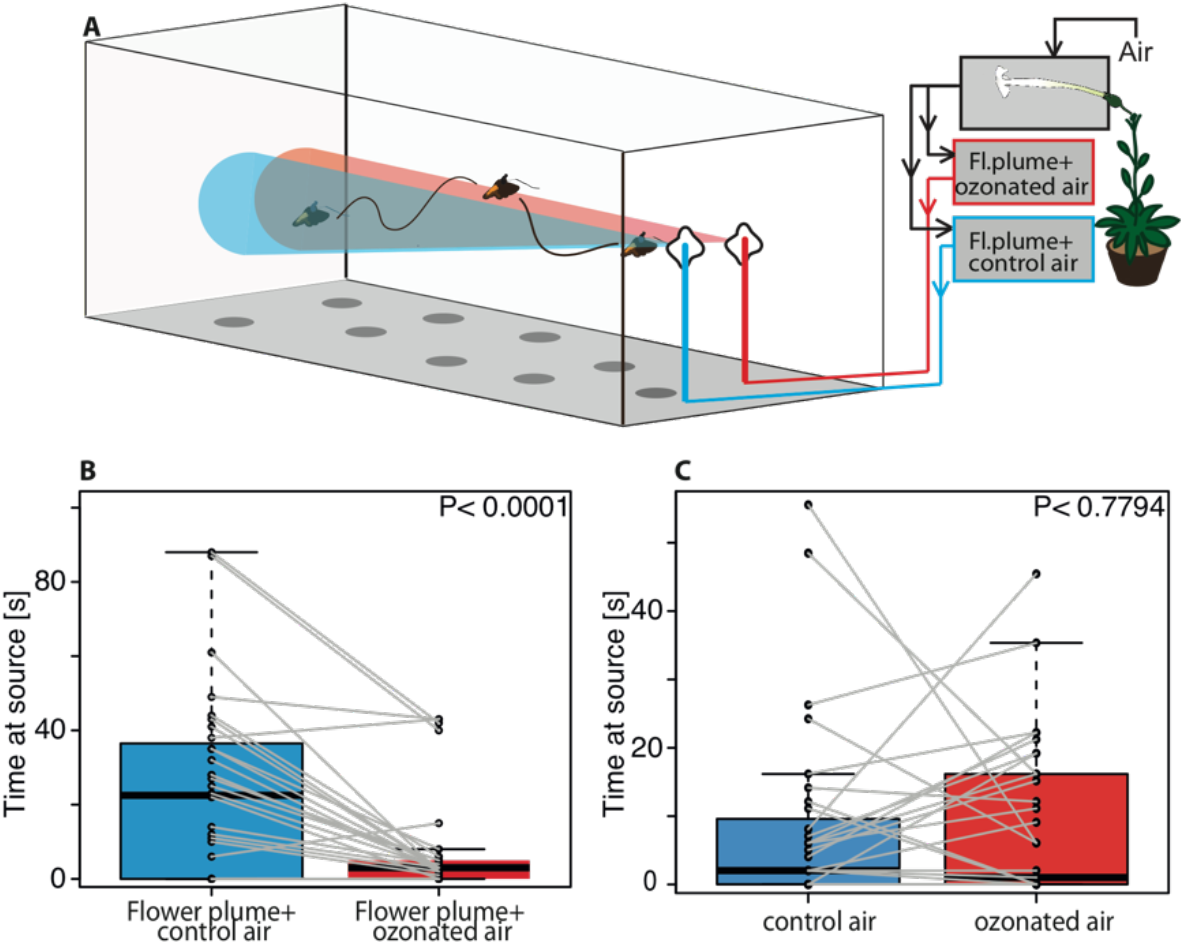
Moths’ innately prefer original to ozonated blends of *N. alata.* A. Schematic of the test for innate preference. B. Time moths spent probing with their proboscis at the artificial flower emitting the original and the ozonated floral blends. (Wilcoxon signed rank test, N=31, p<.05). C. Time moths spent probing with their proboscis at the flower emitting scrubbed air and ozonated air (Wilcoxon signed rank test, N=36, p<.05) (Boxplots, 25% and 75% quartiles and median; whiskers, maximum and minimum values; black circles with connecting lines, paired data of individual moths).

### 2.4 Manduca’s innate response to unaltered vs. ozone-altered floral blends of N. alata

To test the relative, innate preference of moths for the ozone-altered blend compared to the unaltered blend of *N. alata*, individual *M. sexta* flying in the wind tunnel were presented with a choice of two artificial flowers, one emitting the ozone-altered blend, and the other emitting the unaltered blend of *N. alata* (Fig. 2A). Once moths initiated flight in the wind tunnel, they were given four minutes to investigate the two paper flowers, and the amount of time moths spent investigating each ‘flower’ with an extended proboscis was recorded. The two artificial flowers, located side-by-side at the up-wind side of the wind tunnel, were made of light blue paper discs atop metal poles which contained Teflon tubes through which the blends being tested were transported into the wind tunnel. To generate the ozone-altered and unaltered floral blends of *N. alata*, the floral headspace of two *N. alata* flowers housed in a 200ml volume box was split, and half was ozonated by mixing with 120ppbv ozone thorough a series of bottles, while the other half of the floral headspace was mixed only with scrubbed air, following the protocol described above for generating ozone-altered and unaltered floral blends of *N. alata*. The ozonated and pure blends entered the wind tunnel via Teflon tubes at a rate of 0.5 l/min. The side of the wind tunnel from which the ozone-exposed or air-exposed floral blends were emitted was randomly assigned after three moths had been tested.

Next, to establish if any preference for the ozone-altered blend vs. the unaltered blend was driven by an aversion to ozone, the moths were tested for their preference for artificial flowers emitting either 120ppbv ozone or scrubbed air. The flow of 120ppbv ozone and scrubbed air were both emitted from the artifiical flowers at a rate of 0.5l/min. Moths were again given four minutes of flight in the wind tunnel to investigate the two paper ‘flowers’, and the amount of time moths spent at each ‘flower’ was recorded (Fig. 2B-C).

### 2.5 Learning protocol and learned response to ozone-altered floral blend for Manduca sexta

#### 2.5.1 Learning protocol

To test the learning ability of *M. sexta*, we first established a simple three-step olfactory learning procedure using the single odorant Linalool (Fig S1A). In step one, we assessed *M. sexta*’s innate response to Linalool, comparing the attractiveness of a flow of Linalool without obvious visual cues to the attractiveness of a flow of scrubbed air, also without visual cues. Moths were placed at the downwind side of the wind tunnel and presented with a flow of the single floral volatile Linalool produced using 12ul of 10^−2^ Linalool in mineral oil, flowing at a rate of 0.5 l/min from a 250ml airtight bottle, or presented with a 0.5l/min flow of scrubbed air also from a 250ml airtight bottle. Linalool and scrubbed air entered the wind tunnel via the Teflon tubes inside upright metal poles: the structure was the same as used in innate tests, except that visual cues were minimized by removing the artificial flower atop the metal pole. In this initial test without strong visual cues, moths typically did not investigate either odor source, which is in agreement with Bisch-Knaden et al. (2018) who find that Linalool is not innately attractive to foraging *M. sexta*. After a fifteen-minute rest period, the same moth was returned to the wind tunnel for the second step of the learning protocol: training. To train the moths on the odor Linalool, moths were returned to the wind tunnel that now contained a light blue paper ‘flower’ with 12ul of 30% sucrose solution in an epindorf tube at its center, and which emitted the same 0.5l/min of the Linalool odor. Moths were given four minutes to forage on the artificial flower. Those moths that successfully foraged – feeding or attempting to feed for one minute or more, typically until all the sucrose had been consumed – were considered trained. They were removed from the wind tunnel and given another fifteen minute rest interval, after which they were tested in the final step of the learning paradigm: moths were returned to the wind tunnel to repeat their initial air vs. Linalool choice test, again without obvious visual cues. The difference in time spent investigating the Linalool odor relative to air before and after training was used as the metric to assess how well the moth had learned the odor (Fig S1B).

To ensure that moths had learned Linalool and were not merely more responsive to any scent presented after successfully foraging, we switched the training compound, so that instead of being trained and tested on Linalool, moths were trained on 2-Phenyl-ethanol, and then tested to see if their responsiveness to Linalool had improved.(Fig S1C).

#### 2.5.2 Direct learning of ozone-altered floral scents

With a learning system thus established for a single odorant, we proceeded to test *M. sexta’s* ability to learn ozonated floral blends. Following the same three-step learning procedure used for Linalool, we tested *M. sexta’*s intial response to the ozone-altered blend vs. air in the absence of prominent visual cues, and then reassessed that response after the moths had completed their training step, i.e. had foraged on an artificial flower emitting the ozonated floral blend of *N. alata* (Fig. 3B).

**Fig 3.**
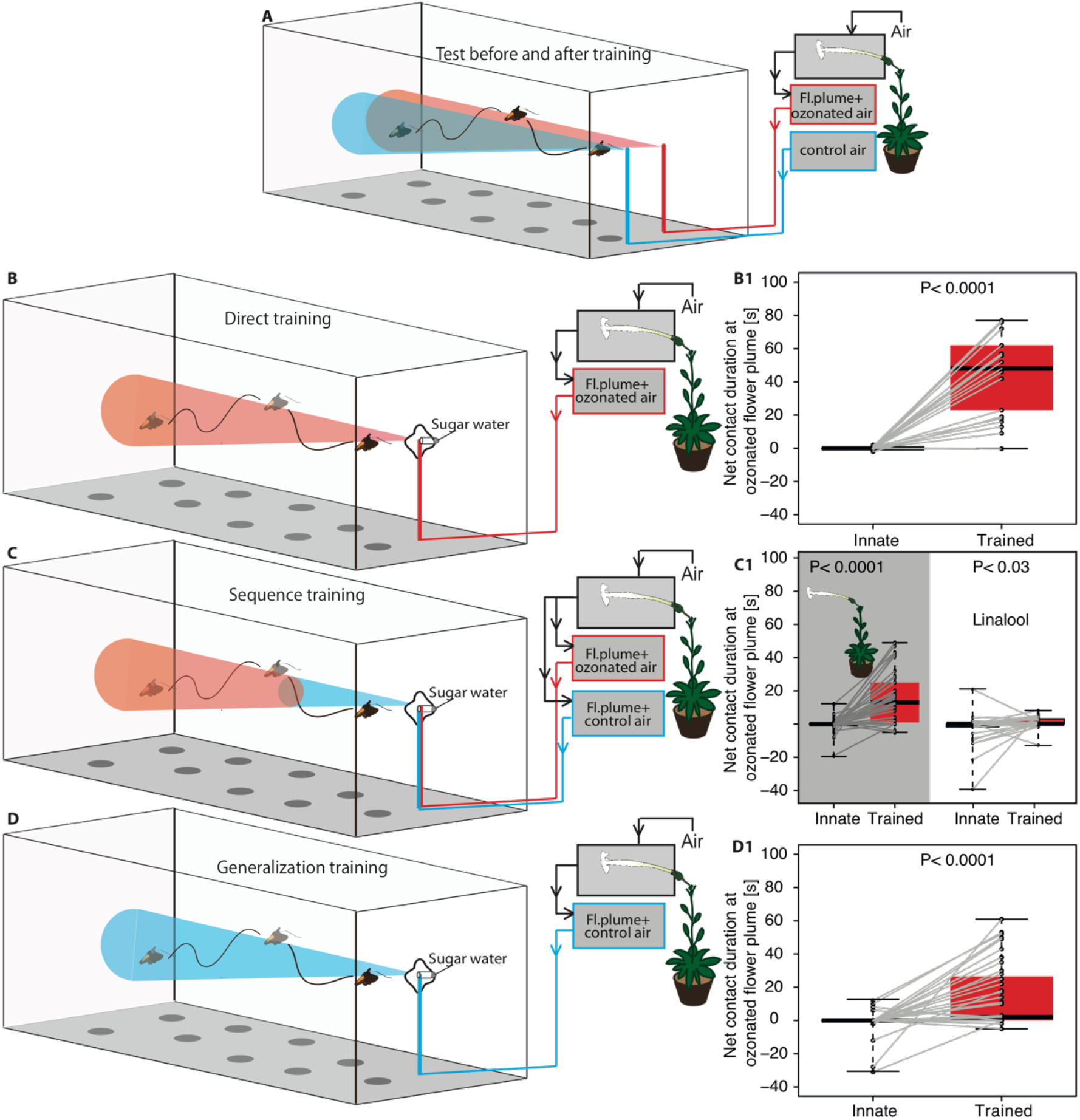
Moths can learn ozonated floral blends. A. Test situation before and after training sessions. B. Direct training, where moths are rewarded with sugar water at an artificial flower emitting an ozonated floral blend. B1. Net contact duration at ozonated floral blend before and after training (Net contact duration; time at ozonated plume minus time at clean air source [s]) (Wilcoxon signed rank test, N=22). C. Sequence training, where moths first follow an ozonated floral blend (red), but then become rewarded with sugar water in the presence of the original blend (blue). C1. Left boxplots, net contact duration at ozonated blend before and after training (N=45), right boxplots, net contact duration at Linalool, when moth in the training situation first had to target a Linalool source, but were rewarded at a flower emitting 2-Phenyl-ethanol) (N=30). D. Generalization training, where moths experience only the original floral blend during the full training situation. D1. Net contact duration at ozonated floral blend before and after training (N=48). (Boxplots, 25% and 75% quartiles and median; whiskers, maximum and minimum values; black circles with connecting lines, paired data of individual moths).

#### 2.5.3 Learning ozonated floral blends as a link to unaltered blends

Close to the flower, floral volatiles have been exposed to ozone for only a short amount of time, resulting in a negligible level of alteration of the floral blend. While moving away from the source through a polluted atmosphere however, the blend chemistry will become more and more altered. Can moths learn that ozonated floral blends lead to rewarding flowers, even though they experience a natural blend at the moment that they receive a nectar reward? There are two potential ways that pollinators could link distinct floral blends (altered and unaltered) to a reward in a natural landscape. They could learn that one stimulus links to a second stimulus, and the second stimulus links to the reward (Hussaini et al 2007). Additionally, they could learn to recognize shared compounds within altered and unaltered floral blends, and treat those unchanging signals as a single continuous stimulus (i.e., treat distinct blends as the same). In order to test these possibilities, we carried out a series of three tests. First, we tested if *M. sexta* could learn to respond positively to an ozonated blend if its experience was followed by a rewarding natural blend. Finding that they could, we used two further tests to try to distinguish the mechanism underlying this response: we tested whether *M. sexta* is capable of linking two distinct stimuli with a reward, and we subsequently tested whether being rewarded at the natural floral blend was sufficient to make the ozonated blend attractive.

We tested if *M. sexta* could learn ozone-altered floral blends if the ozone-altered blend preceded the unaltered blend at the rewarding artificial flower in the training step (Fig. 3C). Using the same initial and final step of the ‘direct learning protocol,’ we modified the training step so that the moth was given one minute to fly towards the artificial flower emitting the ozonated blend of *N. alata* at an increased rate of 1 l/min, until the moth’s extended proboscis was ~8cm away from the source. At this point, we instantaneously switched the scent emitted by the artificial flower from the ozonated floral blend to the unaltered blend using a manually-activated solenoid valve. Moths could then feed at the artificial flower only when it was emitting the unaltered blend. Both the decrease in time allotted for investigation to one minute and the increase in flow of the floral scent from the artificial flower were used to entice the moth to hone in on the floral blend so that the approach was obvious and could be well anticipated by the researcher manually shifting the scent flow at the flower from the ozonated to non-ozonated scent.

Next, we tested whether *M. sexta* could learn to associate a series of two distinctive odors with a reward. We used two single odors instead of an ozonated and unozonated blend to assure that the moth was forming an association for a series of two different cues rather than forming an association for overlapping compounds in the two blends. Following the same two-scent training paradigm, we set up an experiment in which Linalool, a scent not innately attractive to *M. sexta (*Riffell et al 2009; Bisch-Knaden et al 2018), led to a rewarding artificial flower emitting 2-Phenyl-ethanol – an innately attractive scent (Bisch-Knaden et al 2018). The concentration of both Linalool and 2-Phenyl-ethanol was 10ul of 10^−2^ odor in mineral oil from 250ml volume airtight glass bottles.

#### 2.5.4 Learning ozonated floral blends after foraging on unaltered blends

Finally, to test whether learning the ozone-altered blend could be achieved merely from experience with the unaltered one, we ran a third behavior assay where naive moths fed at the original blend and then were tested at the ozonated one (Fig. 3D). Following the same three-step learning procedure described above, moths were first tested for their innate response to the ozonated blend vs. air, were then trained by feeding at an artificial flower emitting only the original blend (i.e., they did not experience the ozonated blend in the training phase), and were finally reassessed for their response to the ozone-altered blend after this training.

## 3 RESULTS

### 3.1 Effects of ozone on the floral headspace of N. alata

Exposing the headspace of *N. alata* flowers to ozone substantially changed the odor profile of the floral blend (Fig. 1 A). The primary difference was a reduction in several compounds in the unaltered blend as well as an increase in decenal. These differences yielded significant separation in the overall blend composition as shown by a principal component analysis (Fig 1B)

### 3.2 M. sexta innate response to ozone-altered floral blend and ozone

After determining that the floral blend of *N. alata* was altered by 120ppbv ozone, we tested the innate preference for *M. sexta* to blends of *N. alata* mixed with air vs. mixed with ozone-enriched air (Fig. 2A). Moths preferred the original blend of *N. alata* to the ozonated one: they spent significantly more time probing the artificial flower emitting the original blend relative to the artificial flower emitting the ozonated blend (Fig. 2B). The decrease in attraction to the ozone-altered blend relative to the original one was not driven by aversion to ozone, as moths in the wind tunnel investigated artificial flowers emitting ozone at 120ppbv versus scrubbed clean air at the same rate (Fig. 2C).

### 3.3 Behavioral learning paradigms for M. sexta foraging on ozone-altered floral blends

Once a learning protocol was developed using a single odor (Fig S1), we tested whether moths could learn the ozone-altered blend using three different learning paradigms. In all three learning paradigms, the initial and final tests – response to the ozone-altered blend vs. air without visual cues – remained the same, while the training step was altered (Fig. 3A-C). Moths learned the ozone-altered blend when the ozone altered one was presented at a rewarding artificial flower, i.e., spent significantly more time at the ozonated blend after the training (direct training, Fig 3B1). Moths were also able to learn the ozone-altered blend when this preceded the unaltered blend at a rewarding artificial flower in a sequence (Fig. 3C1). When tested on their ability to learn a sequence of two single odors rather than blends, however, the moths only slightly increased their response to Linalool when their subsequent reward was paired with 2-Phenylethanol (Fig.3 C1). Finally, even when the moths only experienced the original floral blend (i.e. the ozonated blend being completely absent during the training), they still increased the time they spent investigating the ozonated blend in the subsequent test situation, indiciating an ability to generalize learned odors beyond their innately attractive components (Fig. 3 D1).

## 4 DISCUSSION

### *Ozone alters floral blends and diminishes blend attraction to naïve* M. sexta

Ozone at concentrations currently reached in the Northern Hemisphere can substantially alter the floral blend of *Nicotiana alata* (Fig 1A), a result congruent with earlier modeling and empirical studies (McFrederick et al 2009; Farre-Armengol et al 2016). The *N. alata* blend mixed only with scrubbed air closely resembles the scent profile previously reported for *N. alata* (Raguso et al 2003), while the ozone-exposed blend showed a decrease in some alkenes, including the monoterpene β-Myrcene and the oxygenate monoterpene Linalool, which were reduced via reaction with ozone (Atkinson and Arey 2003). Other compounds in the blend were not affected by ozonation, including Eucalyptol, which has previously been described as non-reactive with ozone (Destaillats et al 2005). Because some compounds react with ozone but not others, the relative ratios of compounds in the blend, and thus the overall blend composition for organisms sensitive to those particular compounds, was altered by ozonation (Fig1B). No new secondary compounds resulting from ozonation of the blend were identified, but a solvent delay in the GCMS protocol prevented low molecular weight compounds such as Acetone, Formaldehyde, or carbon monoxide from being detected. Had these common secondary compounds been detected, the difference between the ozonated and non-ozonated blend would likely have been even greater.

Ozone-alteration of the floral blend was severe enough to reduce *M. sexta’s* innate attraction to *N. alata* (Fig. 2B). This may be due to any of the following changes to the blend composition: a decrease or absence of reactive, innately attractive compounds, a resultant alteration in the ratios of compounds – ratios of odors may be critical for insect recognition of scents (Bruce et al 2005) – or the addition of secondary compounds that may be repellent (Li et al 2016). Additionally, the presence of ozone itself may contribute to this diminished attraction: even though ozone itself was not a deterrent to *M. sexta* in our test trial (Fig 2C), a previous study found that *Apis mellifera* were less responsive to compounds presented in ozone vs. air (Dotterl et al 2016). Regardless of the mechanism behind the loss of attraction, *M. sexta’s* preference for the original blend contributes another case to the body of literature showing that ozone can disrupt innate behavioral responses to attractive compounds (Farre-Armengol et al 2016; Fuentes et al 2013; Li et al 2016).

### Manduca sexta *can learn to be attracted to ozonated floral blends*

Although *M. sexta’s* innate attraction to the floral blend of *N. alata* is disrupted by ozone, associative learning enabled *M. sexta* to find the ozonated blend attractive. The moths readily learned the ozonated blend after they had foraged on a sucrose reward while being exposed simultaneously to the ozonated blend (Fig3B). However, this direct-associative learning mechanism, while providing evidence that ozonated plumes can be learned by the moths, would not be sufficient for *M. sexta* to recognize ozonated blends in the field. In the field, the floral blend in a plume becomes gradually altered by ozone as it moves away from the flower, so that a foraging pollinator would never have the opportunity to forage at a flower while being exposed to the ozonated blend.

In order to learn blends altered by ozone in the field, *M. sexta* would have to learn the ozonated blend decoupled from a reward. Such learning could be accomplished via two mechanisms: a moth could experience the ozonated blend at some distance from the flower, followed by an original blend at the flower, and link the olfactory information from the two blends – i.e. the moths would learn a sequence of ozonated and original blends. Alternatively, feeding at the original blend could reinforce those blend compounds that do not become affected by ozone (e.g. Eucalyptol in Fig. 1A), and a moth could later recognize these unreactive compounds in the ozonated blend, even if these compounds were not innately attractive – i.e., the moths would generalize from original to ozonated blends. *Manduca sexta* that experienced an ozonated floral blend followed immediately by the original blend at a rewarding paper flower, learned to use the ozonated blend as a foraging cue (Fig 3C1 left), which is consistent with learning a sequence of two blends. When, however, we tested this ability to link two odors more explicitly by using the individual compounds Linalool and 2-Phenyl-ethanol as sequential odor cues, *M. sexta* was unable to learn this sequence (Fig 3C1 right). Thus, the learning of a sequence of odors does not seem to account for *M. sexta*’s ability to learn the ozonated blend in the sequence training paradigm.

Instead, moths may have learned by generalizing olfactory information from the original blend to the ozonated one. Indeed, *Manduca sexta* did not need to experience the ozonated blend in the training session at all to later become attracted to it: merely feeding at an artificial flower emitting the original blend resulted in a subsequent increased attraction to the ozonated one (Fig. 3D1). It is apparent that *M. sexta* has learned to recognize identical or structurally similar compounds between the ozonated and original blend, rather than linking the sequence of these blends when approaching the flower. These reinforced compounds could either be those parts of the blend that are not-reactive with ozone (such as common flower odors like Benzaldehyde and Eucalyptol, Raguso et al 2003), or could be compounds in the ozonated blend that are chemically similar (Daly et al 2001) to compounds in the original blend that the moth experienced. Chemically similar compounds may activate the same set of olfactory receptors (Shields and Hildebrand 2001, Bisch-Knaden et al. 2018), and a moth may associate a reward with the activation of that olfactory receptor, regardless of what compound was responsible for its activation. *Manduca sexta* may thus recognize an ozonated blend after foraging on an original one without needing to directly associate non-reactive compounds with rewards. Regardless of which mechanism is being used, this outcome represents a potentially robust flexibility in foraging on preferred host plants in a polluted environment.

Although learning enables *M. sexta* to recognize an ozonated blend in a manner that it may employ in the field, learning as a means of mitigating effects of anthropogenically-induced blend perturbation is not without limits. To begin, ozone is just one air pollutant that has increased as a result of anthropogenically driven fossil fuel combustion: nitrate radical, which peaks at night, has also risen due to increased nitrogen oxides in the atmosphere. In this study, ozone was used as a proxy for nitrate radical, but while both ozone and nitrate radical oxidize alkenes, they do so in a different manner, and nitrate radical can additionally abstract hydrogens from C-H and even O-H bonds (Atkinson and Arey 2003), making it a stronger oxidant than ozone, although it is also less abundant in the troposphere. Furthermore, to associatively learn a floral blend, an insect must first locate a host plant; if floral blends are degraded at long distances this can only be accomplished when an insect comes by chance within either close range or sight of a flower, or if it relies on floral compounds that are not reactive with ozone. An insect’s ability to both recognize and learn olfactory cues, however, is highly species-specific and restricted by the insect’s specific neuro-physiological makeup that is determined by its co-evolutionary history with given host plants. Specialist insects feed on just a few taxonomically related plant species, and naïve specialists may rely on unqiue and taxonomically-distinctive floral volatiles to ensure they can find their host plants (Brandt et al 2017; Schaffler et al 2015; Burger et al 2010; Milet-Pinheiro et al 2012). Thus, a specialist insect’s foraging success in polluted environments may depend on the ozone-reactivity of the few floral compounds that it innately relies on, or the similarity of ozone-induced breakdown compounds to these parent floral volatiles. Furthermore, the ability of specialist pollinators to associate new odors with rewards has not been well assessed, and their ability to use olfactory cues in ozone-rich environments may thus depend entirely on whether or not the few odors they use react with ozone or other oxidants.

In summary, learning can provide one means by which pollinators can still use oxidized floral blends, but the importance of learning as a means of coping with polluted blends in the field remains to be tested. Learning capabilities are likely to be highly variable among species, and, hence, elevated tropospheric oxidants still pose a potentially serious threat to foraging pollinators. Multiple studies have reported declines in insect abundance and pollinator health in regions across the globe (Potts et al 2010; Kluser et al 2007; Fox et al 2013; Hallman et al 2018), and various anthropogenic drivers have been implicated in this decline. Anthropogenically elevated air pollution could be another stressor contributing to overall global insect declines. Future work is needed to assess the real threat of oxidants on foraging insects, and such work must consider pollinators as agents capable of plastic behavioral responses in the field.

## Supplementary material

To test the learning ability of *M. sexta*, we first established a simple three-step olfactory learning procedure for *M. sexta* using the single odorant Linalool (Fig 1A).

**Fig S1.**
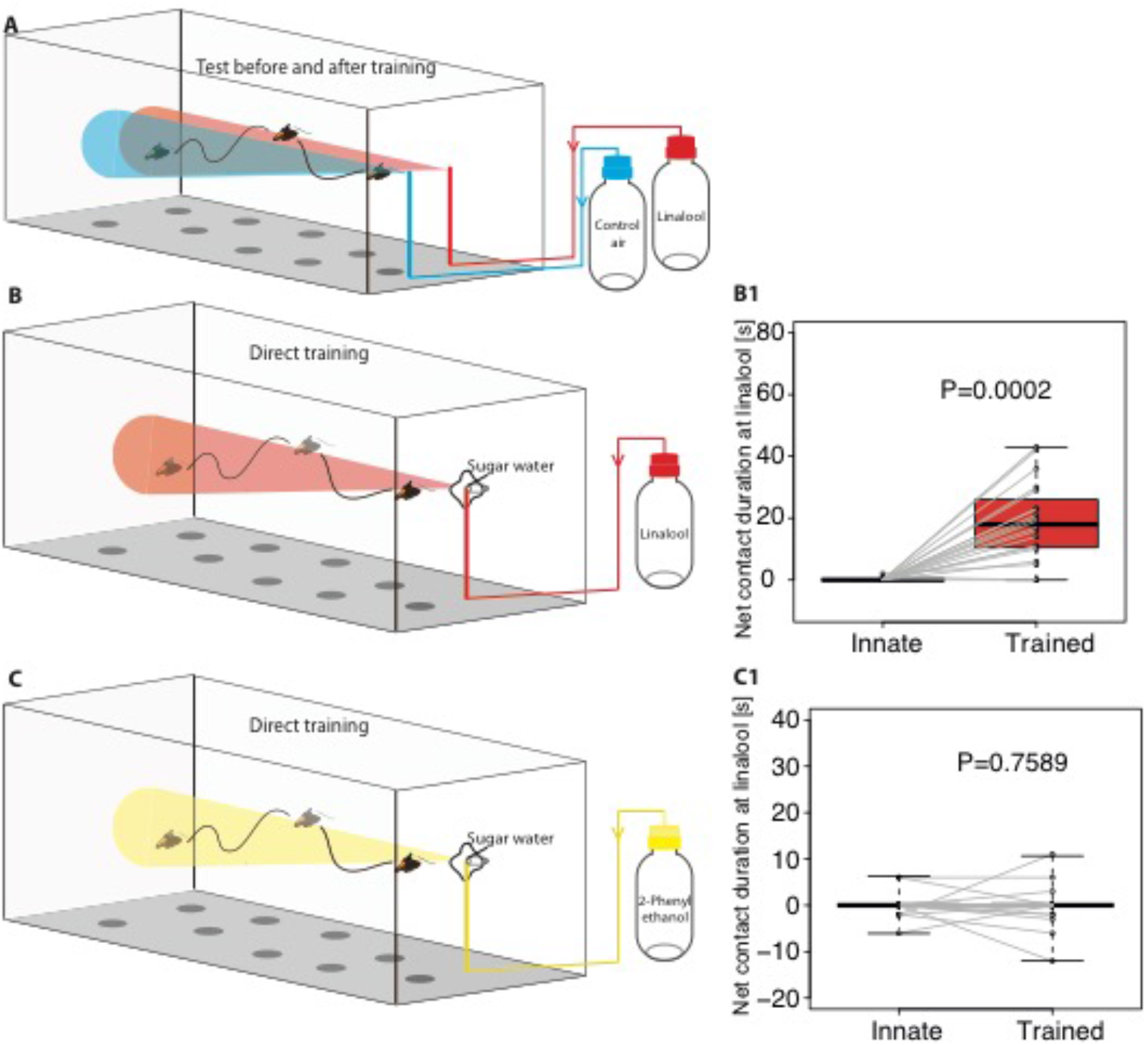
Moths can learn to associate an odor with sugar reward. A. Test situation before and after training sessions. B. Training, where moths are rewarded with sugar water at an artificial flower emitting Linalool. B1. Net contact duration at Linalool source before and after training (Net contact duration; time at Linalool source minus time at control air source [s]) (Wilcoxon signed rank test, N=20). C. Training, where moths are rewarded with sugar water at an artificial flower emitting a 2-Phenyl ethanol plume. C1. Left boxplots, net contact duration at Linalool source before and after training with 2-Phenyl ethanol (N=10).

